# Engineering a Bright Near-Infrared Fluorescent Protein by Screening a Comprehensive Phenotypic Landscape

**DOI:** 10.1101/2025.11.25.690577

**Authors:** Devon L Kulhanek, Qiyao Wei, Jared Head, Jessica Hellinger, Zach Jansen, Andrew Gilmour, Thomas Segall-Shapiro, Jennifer S. Brodbelt, Syed Muhammad Usama, Ross Thyer

## Abstract

Near-infrared fluorescent proteins (niRFPs) are valuable markers for tracking cellular phenomena as they offer greater imaging depth, lower background, and minimal invasiveness relative to other fluorescent probes. The small ultra red fluorescent protein (smURFP) is the brightest niRFP currently reported and has been the subject of several mutagenesis studies to improve its biophysical characteristics. Here, we demonstrate a systematic approach to exploring the mutational landscape of smURFP using a comprehensive deep mutational scanning (DMS) library to identify novel smURFP mutations which confer greater *in vivo* fluorescence. By observing changes in relative abundance between naïve and fluorescence sorted populations, we provide analysis of the enrichment of all possible single-codon substitutions, insertions, and deletions for smURFP. Enriched populations yielded a series of seven single-codon substitutions which confer a three-fold increase in *in vivo* fluorescence in *E. coli* when combined. Finally, we assess the potential underlying mechanisms for increased fluorescence by characterizing the biophysical and photophysical properties of the mutant niRFP sequences. We confirm that two of the derived smURFP sequences have a higher molecular brightness than wild type and yield the brightest niRFP reported.

## Introduction

Near-infrared fluorescent proteins (niRFPs) with peak emission wavelengths over 650 nm are of increasing interest for studies on intracellular phenomena and pathogenesis because of their non-invasiveness and simple subcellular localization relative to synthetic probes. They also take advantage of the deep penetration and low autofluorescence of near-infrared light in both biological tissues and microbial cultures^1–6^. However, most niRFPs suffer from limited molecular brightness due to the low energies of absorbed and emitted photons and are phycobiliprotein derivatives with complicated and inefficient post-translational maturation relative to conventional beta-barrel fluorescent proteins because they require incorporation of conjugated tetrapyrrolic co-factors which function as the chromophore^7–10^. SmURFP is a dimeric niRFP derived from a cyanobacterial phycobiliprotein with absorption and emission maxima at 642 nm and 672 nm, respectively, and it has the highest molecular brightness of any niRFP currently reported^11–16^. To develop an niRFP suitable for tissue imaging, iterative engineering via random mutagenesis, corresponding to ∼15% of the total sequence, was required to make smURFP capable of autocatalytically ligating the non-native chromophore biliverdin (BV) which is generally abundant in mammalian cells. While it has many excellent properties like photostability and mammalian expression efficiency comparable to conventional fluorescent proteins like sfGFP and mCherry, both its unnatural evolutionary trajectory and the limited range of amino acids accessed by error-prone PCR during prior engineering suggest a high potential to find beneficial mutations to improve photophysical properties, chromophorylation behavior, or *in vivo* stability^17^.

Here, we perform a systematic and comprehensive assessment of the phenotypic landscape of nearly all single-codon mutations for smURFP by utilizing DMS to capture nearly all possible single-codon substitution, insertion, and deletion in a single population. We use fluorescence assisted cell sorting (FACS) to observe the phenotypic distribution of the library and isolate highly fluorescent subpopulations which were further subjected to high-throughput screening to identify beneficial single mutations. Screening yielded a series of seven single-codon substitutions which when combined confer a three-fold increase in fluorescence in living *E. coli* cells over the parental smURFP protein. Characterization of the protein both *in vivo* and *in vitro* suggest that changes in smURFP maturation are a primary cause for the increased signal. We also measure the extinction coefficient and quantum yield of each variant in the series of seven iteratively added substitutions and report that two variants have a higher molecular brightness than wild type and represent the brightest niRFPs observed to date.

## Results

*E. coli* does not natively synthesize the BV cofactor required for smURFP maturation, nor is the molecule membrane permeable, and thus requires dedicated biosynthesis from the bacterial heme supply^18^. In addition, evidence from the original engineering work suggests that complete BV ligation quenches fluorescence and that half chromophore occupancy results in maximal fluorescence^12^. Based on this information we hypothesized that smURFP fluorescence would be highly sensitive to the changes in expression of its own sequence as well as the *Synechocystis* Heme Oxygenase-1 (synHO) needed to convert heme directly to BV, and that these two proteins would require careful titration to maintain the equilibrium between chromophore synthesis and ligation within a favorable regime. To assess the impact of smURFP expression, we constructed a plasmid for expression of wild-type smURFP fused to a C-terminal His-tag under the control of P_LtetO_ and co-expressing the cognate TetR transcriptional repressor. The heme oxygenase was expressed from a constitutive promoter on a second plasmid (Figure S1a). This two-plasmid genetic circuit was fine-tuned by titrating promoter and bicistronic design ribosome binding sites (BCD) of different strengths. We evaluated the relative fluorescence outcomes *in vivo* with varied concentrations of anhydrous tetracycline (aTc) to confirm the expected inducible expression of mature smURFP (Figure S2a)^19^. Strong constitutive expression of synHO was lethal or dramatic growth inhibition, potentially due to heme depletion, and we proceeded with the highest expression that did not appear to affect cell fitness. A moderate strength BCD was selected for smURFP expression to achieve high end-point signal without elevated background due to leaky expression.

### Deep Mutational Scanning Library Preparation

The optimized smURFP expression plasmid served as the template for preparation of a DMS library spanning residues K2 through F134 and encoding all possible single-codon substitutions, insertions, and deletions with two exceptions: i) no cysteine substitutions or insertions at any position which may interfere with BV ligation and ii) no substitution or deletion of C52, the wild-type BV binding residue. The library, totaling 5148 unique sequences, was synthesized as five sub-libraries covering different segments of smURFP using a modular Golden Gate assembly scheme to combine dsDNA fragments amplified from an oligonucleotide pool with clonal backbone fragments (see Methods for details) (Figure 1a). Coverage analysis of around 180,000 valid reads from Illumina HiSeq identified 5060 or ∼98% of expected library members were present with minimal bias in abundance apparent between sub-libraries segments (Figure 1b). Additionally, the naïve smURFP library was functionally validated using flow cytometry to visualize the phenotypic distribution of *E. coli* populations co-expressing synHO and either wild-type smURFP or a non-fluorescent C52S mutant which cannot ligate BV, revealing two distinct fluorescence peaks with about a 16-fold difference in fluorescence intensity (Figure 2a). We selected a fluorescence intensity of 200 (a.u.) between these peaks as the threshold for non-fluorescent events, where 100% of C52S events and only ∼8% of wild-type events are under the threshold. The sizeable nonfluorescent subpopulation of cells expressing clonal wild-type smURFP may be due to inherent variance in bacterial cell phenotype or mild synHO toxicity resulting in silencing or genetic instability of the plasmid. Expression of the naïve library under identical conditions yielded a bimodal distribution aligning with the control peaks where approximately 64% of events are under the non-fluorescence threshold (Figure 2a).

**Figure 1:**
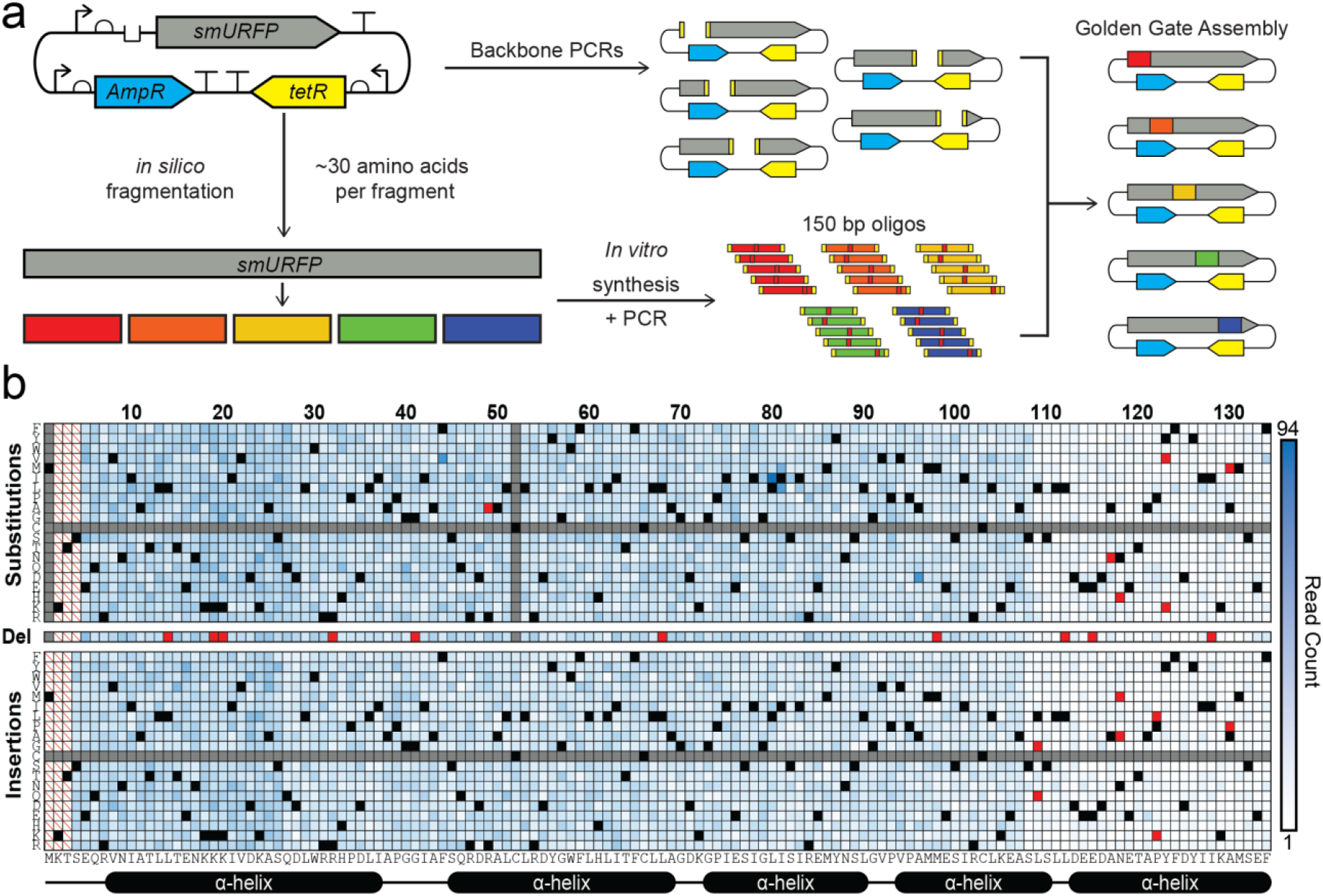
Construction and validation of smURFP deep mutational scanning library. (a) Schematic of the DMS library construction workflow. (b) Heat map showing the direct read count coverage of all possible mutations in the collective naïve library with highlighted wildtype positions (black), unincluded mutations (gray), and unobserved expected sequences (red), and positions locked during amplification (white with red slash).

**Figure 2:**
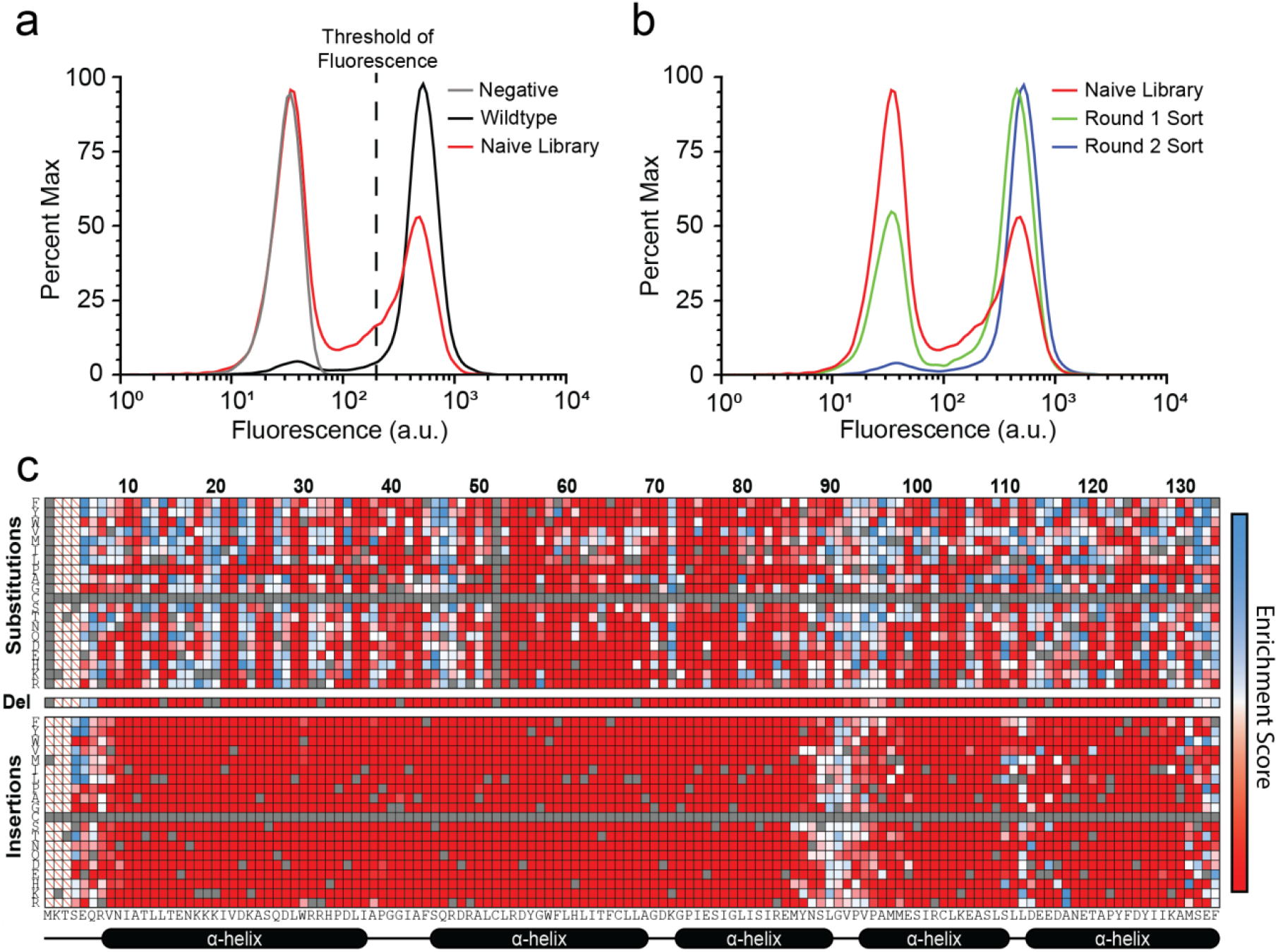
Characterization of library enrichment. (a) Flow cytometry plot showing wildtype (black) and inactive control (gray) populations against the bimodal naïve library phenotype distribution (red). (b) Flow cytometry plot showing the shift of the library phenotype distribution towards primarily active variants after the first (green) and second (blue) top 5% FACS selection. (c) Heat map showing the change in relative abundance of each position/mutation after FACS selection with combinations either enriched (blue), neutral (white), or de-enriched (red). Also highlighting unobserved and wildtype positions (gray) and positions locked during amplification (white with red slash).

### FACS enabled elimination of inactive variants

We used FACS to eliminate deleterious mutations from the naïve library by isolating and subcloning fluorescent variants over two sequential rounds of sorting. In the first round, we performed sorting with gates set to capture the brightest 5% and 2% of events, as well as an “active” sort for any event above the 200 a.u. threshold. All three populations were subcloned and retransformed, and the more stringent 5% and 2% enrichments decreased the rate of non-fluorescent events to ∼40% while the less stringent “active” enrichment resulted in a more modest shift. The population representing the brightest 5% of events from the first round was subject to additional sorting for the brightest 10% and 5%, which both yielded subcloned populations with non-fluorescent events at a similar frequency to the clonal wild-type control (Figure 2b). Around 750,000 valid Illumina HiSeq reads were collected for the subcloned population derived from the two sequential 5% FACS sorts, and coverage analysis showed that 2721 of the 5060 originally captured sequences were eliminated during enrichment, mostly corresponding to deletions and insertions (Figure S3). We applied the formulas below to calculate an enrichment score (ε) for each possible mutation, normalizing both the read count per high-throughput sequencing (HTS) run and the abundance of wild type (A_wt_) in each population (Figure 2c)^20^. The value of one represents a neutral enrichment score, and we observed a range from zero (eliminated) to 18.4 with wild type slightly de-enriched during FACS at a score of 0.84.

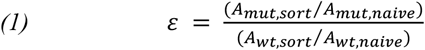

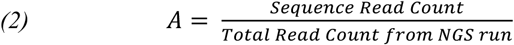

### Screening for beneficial mutations

The naïve and FACS-enriched plasmid pools were transformed into synHO+ cells, and colonies were randomly screened in singlicate to identify mutants producing greater fluorescence than wild-type in bacterial culture. Plasmid DNA was purified from 124 clonal isolates that outperformed wild type in singlicate, retransformed into synHO+ cells, and re-phenotyped in biological and technical triplicate to generate high confidence fluorescence measurements. We subcloned the entire smURFP coding sequence from 29 validated clones and re-phenotyped in replicate, eliminating eight false positives caused by cryptic or unobserved mutations outside of the smURFP sequence. From this process, we compiled a set of 21 unique mutations, including 17 substitutions, three insertions, and one deletion which exceeded wild-type fluorescence in living cells by more than 3% of end-point signal (Figure 3a; Table S1). In agreement with the flow-cytometry results, the naïve library mostly yielded mutants with little to no fluorescence, and most candidate mutations were isolated from the FACS-enriched population (Figure S4).

**Figure 3:**
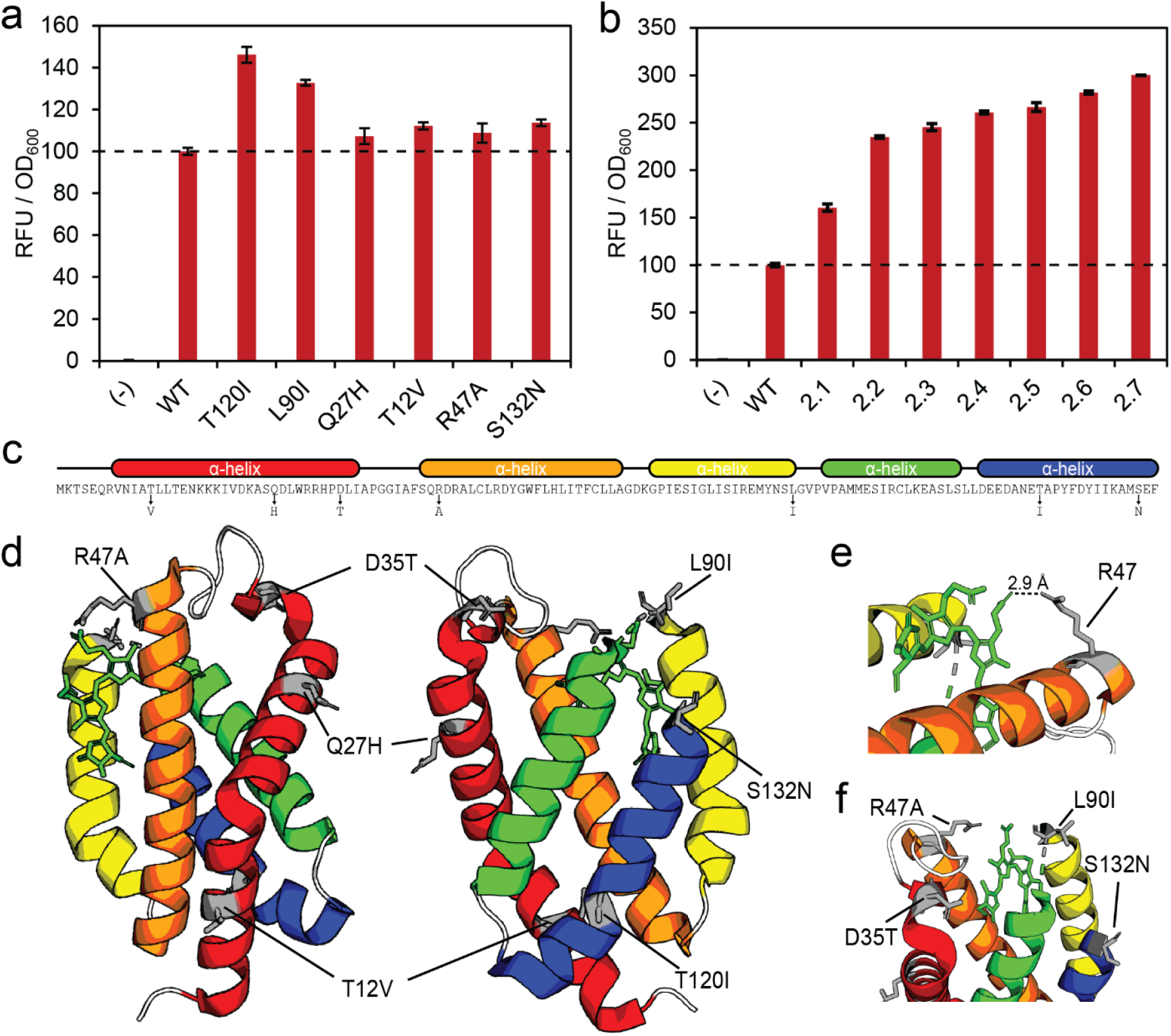
Identification of select mutations. (a) Relative fluorescence plot showing the comparative performance of the single mutation variants screened from the DMS library which were chosen for the combined mutation variants. (b) Relative fluorescence plot showing incremental improvement as mutations are stacked in combination. (c) Substitution locations relative to sequence and color coded α-helices map. (d) Substitution locations relative to full smURFP monomer structure at two different angles. (e) Highlight of the proximity of R47 to the BV chromophore. (f) Highlight of the cluster of substitutions around the BV binding pocket and unstructured loop at L90.

The isolated mutations were not evenly distributed across the coding sequence (Table S1) with nine of the candidate mutations appearing in a dense cluster between S89 and P93 within an unstructured loop, while R47A is the only mutation isolated from the FACS-enriched library between position D28 and I81 (encompassing all of sub-library 2 and 3)^21^. To investigate the possibility that mutations from this region were under-represented or under-performing due to population biases not directly related to smURFP phenotype, we performed additional screening on the two sub-libraries using their naïve plasmid pools but again failed to identify any beneficial mutations. It is worth noting that average HTS read count in the naïve library population for sub-library five, spanning L109 to F134, was 3-fold less abundant than observed for the other sub-libraries, yet mutations from this section were well represented among enriched mutations, suggesting that any beneficial mutations between residues 28 and 81 would have been observed in the whole-population screening as well.

This set of phenotypically validated single-codon mutations were systematically combined by iteratively addition and phenotyping, selecting the best performing combination of mutations as the template for the subsequent round. The single most beneficial mutant was T120I, which was then paired with eight of the other high performing single codon substitutions revealing a nearly two-fold improvement in fluorescence when paired with L90I. This process was repeated over another four rounds to sequentially incorporate T12V, Q27H, R47A, and S132N, and names for each combination based on most recently added mutation are noted in Table 1. The relative end-point fluorescence of this series of variants yielded incremental improvements for mutations three through six, reaching a 2.8-fold increase over wild type (Figure 3b-c).

**Table 1:**
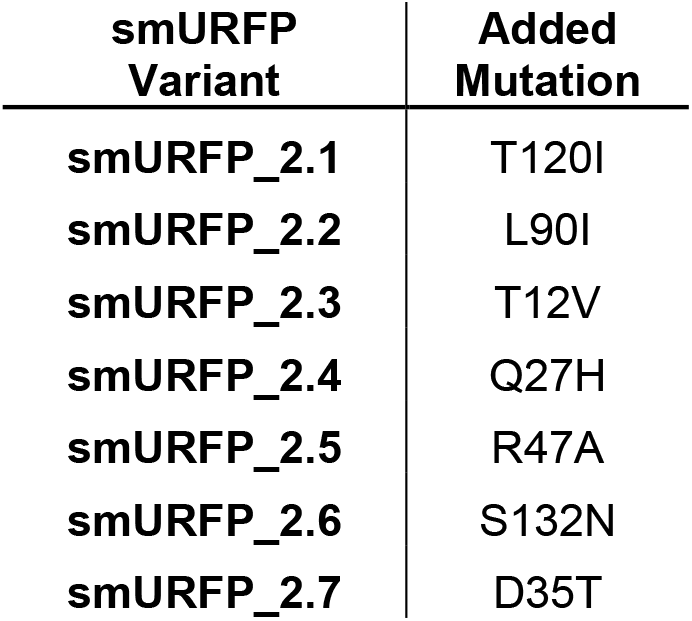
Variant names for iteratively combined amino acid substitutions in order of addition.

Following combinatorial assembly of empirically validated mutations, we also investigated limited site saturation mutagenesis at residues identified from the mutation enrichment scores. While most residues were broadly intolerant of mutation, a subset of residues were highly mutable and yielded many enriched variants. We selected four of these tolerant residues (D23, Q27, D35, and Y126) and designed degenerate oligonucleotide primers to selectively introduce all of the enriched substitutions identified from sequencing analysis into the smURFP_2.6 scaffold. While randomization of D23, Q27, and Y126 did not yield any improvements, D35T increased end-point fluorescence by 13% over smURFP_2.6.

### Characterization of smURFP mutants

The coding sequences for smURFP wild-type, variants 2.1-2.7, and L90I were cloned into a pET plasmid in a bicistronic configuration with synHO under the control of the strong P_T7_ promoter (Figure S1b), and smURFP variants purified from this context were spectrally characterized to determine extinction coefficient (EC) and quantum yield (QY) (Table 2). The absorbance spectrum of mature smURFP features a characteristic peak at 391 nm corresponding to the Soret band of BV which has a constant EC whether in a free or protein-bound state (Figure 4a)^22,23^. Free BV is removed during purification and dialysis; thus, the measured BV concentration corresponds to the concentration of mature chromophores contributing to fluorescence. The EC for fluorescence was determined from the Q-band absorbance peak at 642 nm relative to the calculated chromophore concentration (Figure S6). QY was determined by reference method to a Cy5 derivative, 3,3′-Diethylthiadicarbocyanine iodide, with similar spectral properties using excitation at 600 nm and emission collected between 610 – 800 nm^24^. All variants maintained the same absorption and emission maxima as wild-type smURFP, indicating no changes to the covalent attachment site on BV or the hydrogen bonding environment that affect the conjugated system^25^. In contrast, changes in EC, QY, and overall molecular brightness do not follow a clear trend as mutations are added from smURFP_2.1 to smURFP_ 2.7 despite clearly improving fluorescence in living cells, suggesting that these mutations are not primarily improving the photophysical behavior of smURFP. It is worth noting that smURFP_2.7 had abnormally poor solubility and low purification yields relative to all other variants, and the soluble fraction may not represent the overall chromophorylation state of the protein in cells and in turn the observed spectral properties. For example, a greater abundance of dimers with two ligated BV would cause significant fluorescence quenching and suppress QY^12^. Finally, smURFP_2.1 and smURFP_2.5 have 11% and 38% higher molecular brightness than wild type, respectively, and represent the brightest niRFPs reported to date.

**Table 2:** Summary of the photophysical properties of smURFP mutants in 1x PBS (pH 7.4) including extinction coefficient, quantum yield, and the resulting molecular brightness. Quantum yield was determined in reference to 3,3’-Diethylthiadicarbocyanine iodide in methanol (QY = 0.33)^24^.

| Sample | $\epsilon$ (M<sup>-1</sup> cm<sup>-1</sup>) | $\Phi_f$ (%) | Brightness |
| --- | --- | --- | --- |
| Wildtype | 1.4 x 10<sup>5</sup> | 26 | 37,000 |
| T120I | 1.6 x 10<sup>5</sup> | 26 | 41,000 |
| L90I | 1.5 x 10<sup>5</sup> | 17 | 26,000 |
| smURFP_2.2 | 1.6 x 10<sup>5</sup> | 21 | 34,000 |
| smURFP_2.3 | 1.2 x 10<sup>5</sup> | 23 | 28,000 |
| smURFP_2.4 | 1.3 x 10<sup>5</sup> | 25 | 33,000 |
| smURFP_2.5 | 1.6 x 10<sup>5</sup> | 32 | 51,000 |
| smURFP_2.6 | 1.3 x 10<sup>5</sup> | 29 | 38,000 |
| smURFP_2.7 | 0.82 x 10<sup>5</sup> | 19 | 15,000 |

**Figure 4:**
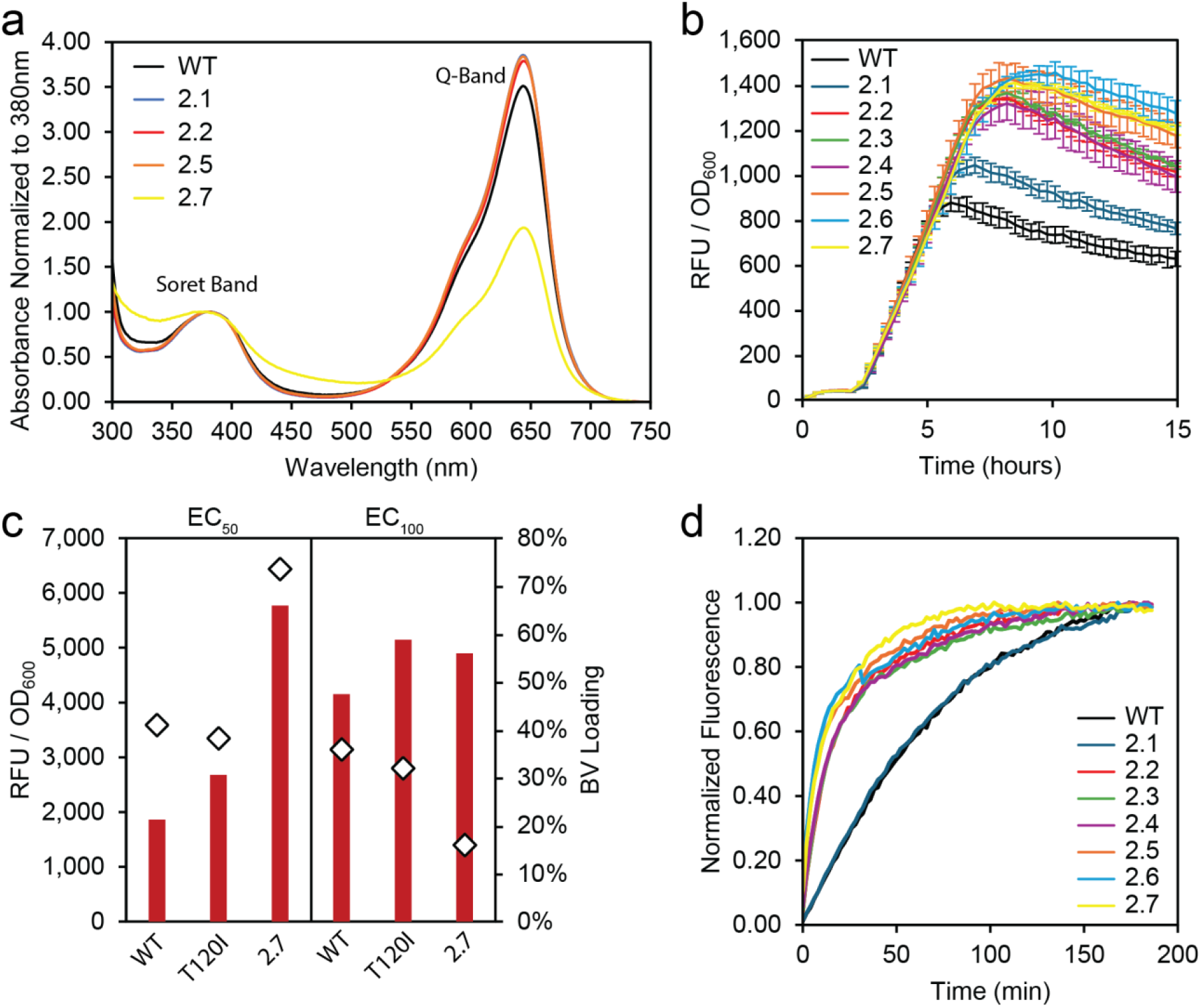
Biophysical characterization of smURFP mutants. (a) *In vitro* absorption spectra for several smURFP variants showing differences in the ratio of Soret and Q-band peaks. (b) Continuous measurement of RFU/OD_600_ for smURFP wild-type and 2.1 – 2.7 in the TetR-controlled expression context induced with 6 ng aTc/ mL. Measurements taken in biological triplicate and technical duplicate, and data represents the mean fluorescence normalized to cell density ± standard deviation. (c) End-point RFU / OD_600_ (red bars) of smURFP wildtype, T120I, and 2.7 from 200 mL baffle flask cultures induced with 6 ng aTc/ mL or 12 ng aTc/ mL and the corresponding BV ligation loading amount (diamonds) among smURFP monomers determined by mass spectrometry of purified protein. (d) Plot of kinetic rate of BV covalent ligation measured *in vitro* at 25 °C.

When assessing the performance of smURFP_2.1-2.7 *in vivo*, the relative end-point fluorescence among variants was found to be highly sensitive to assay parameters with higher expression mildly reducing differentiation and short culture durations (∼6 hours) largely eliminating differentiation (Figure S7a). Furthermore, when expressed in the bicistronic context where synHO expression and resulting BV production is expected to be much higher than in the circuit with constitutive synHO expression there is no significant difference in endpoint fluorescence between variants at all (Figure S7b). sfGFP matures rapidly without cofactor requirements and can be used as a reference for the dynamics of inducible protein expression *in vivo* when RFU/OD_600_ is measured continuously in a microtiter plate. Comparing the development of fluorescence using this method reveals that wild-type smURFP ceases maturation soon after protein expression stagnates while fluorescence for smURFP_2.7 continues to increase for several hours (Figure S8a-c)^26,27^. When wild-type smURFP and the series of variants are subject to weaker induction near the EC_50_, fluorescence accumulates at a nearly identical rate until four hours post-induction when protein synthesis slows, and the series diverges to roughly the expected rank order seen in end-point fluorescence assays as variants continue to accumulate signal for longer periods (Figure 4a). The same dynamics were observed with aTc concentrations near the EC_100_ except the divergence point shifted to around ten hours (Figure S8d-f). Thus, a key contributor to the increased fluorescence of the smURFP variants in living bacteria appears to be an extended maturation phase that persists for several hours after the wild-type smURFP reaches maximum fluorescence, prior to a clearly defined signal dilution phase as the culture turbidity increases.

This trend in RFU/OD_600_ was maintained at larger culture scales. When smURFP wild-type, smURFP_2.1, and smURFP_2.7 were expressed in the TetR-regulated plasmid context in 200 mL cultures in baffled flasks at both the EC_50_ and EC_100_ of induction, lower expression resulted in greater difference between the variants while higher expression suppressed the observed fold-change (Figure 4c). The amount of covalent BV attachment per smURFP monomer (i.e., “loading”) was determined by mass spectrometry for each of the three proteins for two expression conditions (6 ng aTc/mL and 12 ng aTc/mL) (Figure 4c; Figure S9a-b)^12^. It is important to note that smURFP_2.7 consistently yielded a much smaller quantity of soluble protein than all other variants, although the total protein in cell lysate was similar. We observed similar chromophorylation amounts for wild type and smURFP_2.1 with 41% and 38%, respectively, at lower induction and a slight decrease to 36% and 32% at higher expression, as expected given no commensurate increase in BV concentration. In contrast, smURFP 2.7 was found to have a 74% BV attachment at low expression but only 16% at high expression.

In the cell, smURFP apo-protein rapidly dimerizes after synthesis and maturation of the fluorescent chromophore proceeds by a two-step mechanism: i) BV caging within the binding pocket, and ii) covalent attachment to C52 via a thioether crosslink^10,12^. Caging alone does not yield an active chromophore, and Rodriguez, *et al*. confirmed that the rate of initial binding is very fast compared to the rate of covalent attachment by measuring *in vitro* fluorescence maturation of smURFP apo-protein mixed with free BV under different reducing conditions to determine kinetic rates^12,28,29^. When fitted to an exponential rate law, the reaction half-life (T_1/2_) for wild-type smURFP was determined to be of 49 min. We prepared non-fluorescent apo-protein for several of the smURFP variants derived from the DMS library and performed similar *in vitro* kinetic measurements. Fitting to the same exponential rate law determined T_1/2_ for smURFP wildtype and T120I as ∼60 minutes under our experimental conditions and T_1/2_ for all variants carrying L90I as ∼20 minutes, indicating that the L90I mutation drastically improves the covalent ligation rate of BV once caged in the binding pocket (Figure 4d).

## Discussion

Prior optimization of smURFP has relied on error prone PCR to generate random variation and accepted a constrained view of the mutational landscape to broadly and agnostically search for beneficial mutations. This includes the original development of smURFP from a phycobiliprotein and two subsequent attempts to monomerize the protein either by targeted mutagenesis of residues at the dimerization interface or by computational generation of two additional N-terminal helices to replace the interactions otherwise found at the dimer interface^12,21,30^. Both attempts at monomerization resulted in a substantial loss of fluorescence in living bacteria and only achieved partial recovery after iterative rounds of random mutagenesis. By performing sequence enrichment analysis of a smURFP DMS library subject to FACS screening for retained fluorescence activity, we have captured a comprehensive phenotypic landscape of nearly all single-codon mutations, including insertions and deletions, spanning the entire coding sequence. This approach allowed us to both assess broad phenotypic trends in the sequence space and efficiently sample the enriched population to identify dozens of beneficial mutations. Combinations of these mutations yielded new smURFP variants with improved photophysical properties, including smURFP_2.5, the brightest niRFP reported, as well as variants with up to three-fold greater fluorescence in living bacteria.

From just two high-throughput sequencing datasets, we observed the global enrichment of single-codon mutations, which captured both expected biophysical properties and identified several hotspots which proved highly tolerant of mutation (Figure 2c). Consistent with a global mutational landscape, insertions and deletions proved highly deleterious, and proline was poorly tolerated relative to other amino acids^31,32^. Insertions are likely to disrupt local structure at most positions, especially mid-helix, and were only enriched among the six residues at either termini or within the unstructured loops centered on L90 and L111 where the protein is flexible enough to accommodate them. Single-codon deletions are also likely to disrupt the overall structure of the protein and were not enriched at any position. Similarly, structurally constrained residues L51 and R54 – Y56 which flank C52 and form the chromophore binding pocket were intolerant of mutation^21^. In contrast, we can clearly identify the spacing of tolerant positions within the N-terminal helix between residues K20 and G40 where solvent facing residues are broadly tolerant of mutation and interior facing residues are broadly intolerant^33^. Notably, the two most impactful mutations in the final set of seven substitutions, T120I and L90I, were preferentially enriched mutations at their respective residues.

While it can be difficult to ascribe an exact reason for why any single mutation improves performance, the biophysical and photophysical characterization of our smURFP variants hints at a variety of potential mechanisms. The single most impactful mutation, T120I, replaces a polar residue embedded in the core of the protein with a hydrophobic moiety and results in an 11% increase in molecular brightness solely through an increased extinction coefficient (Figure 3f). It is possible that this mutation increases the hydrophobicity of the environment between helices and improves protein stability. The second mutation in the combinatorial series, a conservative L90I substitution, occurs at one of the most mutable locations in the protein and lies immediately adjacent to a loop. Due to the flexibility, this region was not resolved with the rest of the structure^21^. However, many different mutations in this region were isolated as beneficial during screening. From our *in vitro* characterization of covalent chromophore attachment, the rate of BV ligation increased in any combination of mutations where L90I is present, and T_1/2_ is decreased to a third of the wild-type value. Among the other five substitutions characterized in various combinations, specific causes for improved fluorescence were less obvious although we observe that these mutations consistently improve the duration of the protein maturation period *in vivo* (Figure 4a). It is worth noting that four of these mutations were spatially clustered near the entrance of the BV binding pocket including D35T, R47A, L90I, S132N (Figure 3g). The side chain of R47 carries a strong positive charge in close proximity to BV and was replaced with a smaller hydrophobic side chain unable to make direct interactions (Figure 3e).

Many research applications would benefit from brighter niRFP genetic probes that can leverage excitation wavelengths with excellent penetration depths and emission peaks outside of the window of autofluorescence to visualize a wide variety of tissue and other phenomena^34–37^. In addition, the thioether crosslink of smURFP has been adapted to function as a reporter for incorporation of the non-canonical amino acid, selenocysteine, which is likewise capable of tethering the BV through a selenoether linkage^13^. However, niRFPs often have complex maturation pathways and poor photophysical brightness relative to conventional beta-barrel probes like sfGFP^7^. A complex network of interacting residues affect maturation and photophysical behavior, making it difficult to design targeted libraries, but our DMS and global phenotyping strategy offered a systematic solution to this problem. Additionally, we found that expression context strongly affected the relative performance of smURFP variants. The proportion of smURFP apo-protein to available BV in a system strongly affected end-point assay measurements and we found that it was important to control this ratio to achieve meaningful differentiation. We suspect that maximizing over-expression without proportionally greater BV synthesis during screening does not reflect application-relevant conditions^38,39^. Furthermore, isolating protein variants which are photophysically brighter on a per molecule basis is desirable to improve signal sensitivity regardless of expression level; however, separating whether a mutation is affecting expression and maturation versus molecular brightness is not possible from *in vivo* fluorescence alone, and characterizing photophysical properties of mutants requires laborious and low-throughput purification and *in vitro* measurement. Use of time course measurements for *in vivo* fluorescence and other techniques which can capture more aspects of protein maturation will be helpful towards designing screens that discriminate between changes in molecular brightness, chromophore ligation, and protein expression.

## Methods

### Molecular Biology

All plasmids described in this work were constructed using either Gibson Assembly or a previously reported hierarchical, Golden Gate assembly system^19,40^. Gibson Assemblies were performed with a 1:3 ratio of backbone: insert DNA and incubated at 50 °C for 2 hours. Golden Gate assembly reactions were performed by combining 20 fmol of each DNA fragment, T4 DNA ligase, T4 DNA ligase buffer, and the appropriate restriction enzyme. Assemblies were transformed into *E. coli* DH10B or Marionette-Clo using chemical transformation.

### DMS Library Design, Construction, and Validation

An algorithm (SPINE) for generating oligonucleotide tiles to introduce a scanning library of insertion sites to a target gene was previously reported and made publicly available^41^. We modified this python script to output custom oligonucleotides for constructing deep mutational scanning libraries instead (available at www.github.com/Thyerlab/SPINE_Thyer). Briefly, the computational pipeline will: i) accept FASTA files with the gene-of-interest in the entire plasmid or limited upstream and downstream context as input; ii) fragment the target sequence into segments with unique Golden Gate overhang identities and appropriate lengths based-on a user-defined output oligo size; iii) append user-defined type-IIs restriction sites for Golden Gate cloning and adapter sites for amplification to each fragment sequence; iv) generate iterations of each fragment sequence encoding every desired single-codon substitution, insertion, or deletion and excluding the wildtype or stop-codon sequences; v) eliminate duplicate sequences and incidental restriction sites and output the complete oligo pool list as a FASTA file and a CSV file; vi) generate a set of oligonucleotide primer sequences for amplifying a clonal backbone from the library template plasmid with corresponding BsaI cloning sites for each fragment as well as the primers to amplify fragment-specific oligos based on the amplification tags.

The algorithm fragmented the smURFP coding sequence spanning residues K2 to F134 into five segments of 78 to 81 base pairs with BsaI cloning sites and generated 5148 unique 150-mer oligonucleotides encoding the DMS library. Cysteine was excluded as a substitution or insertion at any position, and no modifications were made at cysteine 52 which covalently ligates the chromophore. The single pool of dsDNA oligonucleotides encoding the smURFP DMS library was ordered from Agilent Biosciences.

The sub-libraries for each of the five unique insert fragments were prepared separately to minimize population bias. Insert fragments were amplified from the oligonucleotide pool, and corresponding backbone fragments were amplified from the template plasmid. Golden Gate assemblies with BsaI were performed for each insert-backbone pair, the products were purified and electroporated into *E. coli* DH10B. The total pool of transformants were cultured in TB with 100 µg ampicillin/ mL, and three milliliters of confluent culture were harvested to obtain supercoiled plasmid stocks of each sub-library. Finally, DNA from each sub-library was prepared as an equimolar mixture and electroporated into DH10B, cultured, and harvested to obtain a supercoiled DNA stock of the complete naïve library. Sequence coverage was assessed by amplifying the region from residue E5 to F134 with terminal adapter sequences and submitting the purified amplicon products to commercial HTS service (Illumina HiSeq through Azenta AmpliconEZ). Due to priming site constraints K2, T3, and S4 were fixed as wildtype.

### Fluorescence Assisted Cell Sorting and Flow Cytometry

Samples for FACS (SONY SH800S) were prepared by inoculating five mL M9YE cultures with library populations from glycerol stocks or clonal controls from single colonies and incubating at 37 °C overnight. Confluent cultures were diluted 200-fold into M9 medium supplemented with 2.5 g/L yeast extract (M9YE) and the appropriate antibiotics and incubated at 37 °C for two hours at which point smURFP expression was induced with aTc at the empirically determined EC_50_ and incubated at 37 °C for a further four hours. Cultures were harvested by centrifugation (2000 x *g* and 4 °C), washed twice with three mL of sterile PBS, and finally resuspended in five mL of sterile PBS. Serial dilutions were prepared in sterile PBS at 10x, 100x, and 1000x. Samples for flow cytometry were prepared similarly but using one mL cultures in a 96-well deep-well block. For FACS, populations were roughly sampled to determine appropriate settings to achieve ∼1000 events/ second, and sorting into fresh TB media with appropriate antibiotics was performed until 50,000 events were collected. To subclone and eliminate errant plasmid backbones, the entire smURFP coding sequence was amplified from plasmid DNA recovered from the sorted population and inserted into clonal backbone by Gibson assembly.

### Fluorescence Assays

All fluorescence assays were performed in *E. coli* strain DH10B grown in M9YE unless otherwise specified. Biological triplicates of transformants were selected in 1 mL cultures in 96-well deep-well plates and incubated at 37 °C overnight with 900 rpm agitation on a 1.5 mm orbit. Following overnight growth, confluent cultures were diluted 500-fold into fresh culture medium supplemented with antibiotic(s) in a 96-well deep-well plate and incubated for two hours after which cultures were induced with 6 ng aTc/ mL unless otherwise specified. Cultures were incubated for an additional fourteen hours and then harvested by centrifugation at 3500 x *g* for ten minutes. Cell pellets were resuspended in 1 mL PBS, and 100 µL aliquots were transferred to a 96-well microtiter plate. Fluorescence assays were conducted using a multimode plate reader (Tecan M200 Pro). Absorbance for culture density was measured at 600 nm and fluorescence was measured at λ_ex_ 630 nm and λ_em_ 680 nm. Unless otherwise indicated, all data for fluorescence assays was collected from a minimum of three biological replicates and 3 technical replicates. Error bars represent the standard deviation.

Time course measurements were performed in microtiter plates in a similar manner to block-based assays. Plates were prepared with 87 µL of M9YE with antibiotic(s) in each sample well, inoculated with 3 µL of confluent culture, and incubated at 37 °C with agitation in a Tecan M200 Pro. Measurements were taken for OD_600_ and fluorescence every 20 minutes. 10 µL of fresh media was added containing 10x the desired final concentration of aTc or IPTG after two hours.

### Protein Expression and Purification

Variants of smURFP were expressed for purification from a derivative of pCDFsolo either by itself to yield apo-protein or in a bicistronic arrangement with synHO to yield mature and fluorescent protein (Figure S1a and b). A single transformant was used to inoculate 3 mL of M9YE with 100 µg spectinomycin/mL and 2% (w/v) glucose and incubated with agitation overnight at 37 °C. One mL of the starter culture was used to inoculate a 500 mL baffled flask with 200 mL of M9YE + 100 µg spectinomycin/mL. Cultures were incubated with agitation at 37 °C until reaching an OD_600_ of ∼ 0.6 before smURFP expression was induced with 25 µM IPTG. Each culture was allowed to incubate at 37 °C for another 16 hours before pelleting the entire cell mass for storage at -80 °C until purification. Cells were resuspended in IMAC resuspension buffer (50 mM NaH_2_PO_4_, 100 mM NaCl, 1 mM EDTA, pH 8.0) with 3 mg lysozyme/mL and incubated at 37 °C for 60 minutes followed by further lysis by sonication. Lysate was clarified by centrifugation and the EDTA quenched by addition of MgCl_2_ to 10 mM final concentration. His-tagged smURFP was collect on Ni-NTA resin column and eluted with IMAC elution buffer (50 mM NaH_2_PO_4_, 300 mM NaCl, 300 mM imidazole, pH 8.0). Protein was dialyzed into 1x PBS for storage and characterization using 10,000 MWCO cassettes. Protein concentration was measured by BCA assay.

### Photophysical Characterization

All absorbance spectra were obtained on JASCO V-770 spectrophotometer in 1x PBS (pH 7.4, Gibco). Extinction coefficient (EC) was determined by Beer-Lambert-Bouguer equation through comparison of Q band absorbance (642 nm) to BV peak absorbance (391 nm) according to the method described^23^. The Q band absorbance was kept between 0.1 – 1 nm. The absorbance spectra were measured from 250-750 nm (1 nm step size). Spectra were recorded with synthetic quartz cuvette (ThorLabs; CV10Q35FAE) holder with 10 mm path length.

Fluorescence emission of FPs was measured using FluoroMax from Horiba scientific with 4 nm excitation and emission slit widths, and a 0.1 s integration rate. All FPs were excited at 600 nm, and emission was collected between 610-800 nm. Measurements were carried out at a concentration with an optical density of Q-band less than 0.1 in 1x PBS pH 7.4 (Gibco). Spectra were recorded with synthetic quartz cuvette (ThorLabs; CV10Q35FAE) holder with 10 mm path length.

Fluorescence quantum yield (Φ_f_) was determined relative to 3,3′-Diethylthiadicarbocyanine iodide (Sigma-Aldrich) standard (Φ_f_ = 0.33 in MeOH_2_) for optically dilute solutions (absorbance in the range from 0.01-0.05)^24^. Excitation and emission parameters were matched for the reference dye and FP variants tested. The parameters used to gather the quantum yields were excitation at 600 nm, emission to 610-800 nm, and a slit width of 4 nm. Spectra were recorded with synthetic quartz cuvette (ThorLabs; CV10Q35FAE) holder with 10 mm path length.

The fluorescence quantum yield was calculated using the following formula:

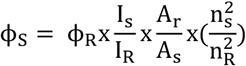

ϕ_S_ = Quantum yield of the sample

ϕ_R_ = Quantum yield of reference

I_s_ = Intregrated fluorescence area of the sample

I_R_ = Integrated fluorescence area of the reference

A_r_ = Absorbance of the reference at the excitation wavelength

A_s_ = Absorbance of the sample at the excitation wavelength.

η_s_ = Refractive index of the sample

η_R_ = Refractive index of the reference

### Mass Spectrometry

SmURFP proteins were diluted to 10 μM and desalted with Amicon 10 kDa MWCO filters into 50:50 methanol:water with 0.1% formic acid. Protein solutions were analyzed on a Thermo Orbitrap Eclipse mass spectrometer (Thermo Fisher, San Jose, CA) using direct infusion nanoelectrospray ionization. Protein solutions were introduced with a Au/Pd-coated borosilicate static tip pulled in-house (OD 1.2 mm, ID 0.69 mm) with an applied voltage of 1.4 kV. Spectra were collected at 240 K resolution with 200 averages, using full profile mode. MS1 spectra were deconvoluted in Xtract in Freestyle with a S/N of 10. Semi-quantitative analysis of loading amounts was accomplished using the relative abundances of smURFP with and without ligated biliverdin in the deconvoluted MS1 spectra. Relative abundances were determined based on the summed peak intensities of all charge states of the proteins (using Xtract in FreeStyle, Thermo Scientific), and it was assumed that the attached biliverdin had minimal impact on the ionization efficiency of the protein. The loading percentage was calculated by dividing the summed intensity of the BV-ligated protein by the summed intensities of the BV-ligated and non-ligated proteins. All measurements were an average of three replicates with an average standard deviation of 1% in the loading amounts based on all replicates.

### *In Vitro* Maturation Assay

*In vitro* kinetics measurements were taken in microtiter plates in a Tecan M200 Pro in 100 uL aliquots at 25 °C or 37 °C. 90 µL of apo-protein solution normalized to 5 µM dimer (10 µM BV binding sites) were quickly mixed with 10 µL of 10x PBS solutions of commercially available BV (Frontier Specialty Chemicals, Cat. #: B655-9) to reach a target concentration of 10 µM and quickly transferred to the plate reader to track fluorescence with excitation at 630 nm and emission at 680 nm. Kinetic parameter fitting for an exponential rate law for generated fluorescence (*F* = 1 − *e*^−*kt*^) was performed to maximize an R^2^ value between the empirical measurements and fitted equation, and *T*_1/2_ = In(2)/*k*_*fit*_.

## Associated Content

## Author Information

### Authors

**Devon Lee Kulhanek –** *Department of Chemical and Biomolecular Engineering, Rice University, Houston, TX*

**Qiyao Wei –** *Department of Bioengineering, Rice University, Houston, TX*

**Jared Head –** *Department of Chemistry, The University of Texas at San Antionio, San Antonio, TX*

**Jessica Hellinger –** *Department of Chemistry, The University of Texas at Austin, Austin, TX*

**Zach Jansen –** *Systems, Synthetic, and Physical Biology Graduate Program, Rice University, Houston, TX*

**Andrew Gilmour –** *Systems, Synthetic, and Physical Biology Graduate Program, Rice University, Houston, TX*

**Thomas Segall-Shapiro –** *Department of Pathology and Genomic Medicine, Houston Methodist Research Institute, Houston, TX*

**Jennifer S. Brodbelt –** *Department of Chemistry, The University of Texas at Austin, Austin, TX*

**Syed Muhammad Usama –** *Department of Chemistry, The University of Texas at San Antionio, San Antonio, TX*

### Notes

The authors declare no competing financial interest.

## Acknowledgements

The authors thank the Welch Foundation for their support (235019 to R.T. and F1155 to J.S.B.). D.L.K. is grateful for the support of the Graduate Research Fellowship Program (National Science Foundation). S.M.U. acknowledges The Welch Foundation (AX-2216-20240404), and The Max and Minnie Tomerlin Voelcker Fund (SAT0004356). J.S.B. acknowledges support from NIH R35GM139658.

