## Supplementary Information for "Engineering a Bright Near-Infrared Fluorescent Protein by Screening a Comprehensive Phenotypic Landscape"

This file contains:

**Supplementary Figure 1** | Schematics of genetic circuits

**Supplementary Figure 2** | aTc induction gradients for the TetR-controlled circuit

**Supplementary Figure 3** | FACS-Enriched library NGS coverage heat map

**Supplementary Figure 4** | Naïve versus enriched library singlicate screening example results

**Supplementary Figure 5** | Absorbance and fluorescence spectra from *in vitro* measurements

**Supplementary Figure 6** | Absorbance measurements for extinction coefficient calculation

**Supplementary Figure 7** | smURFP_2.1-2.7 phenotype under varied assay conditions

**Supplementary Figure 8** | Continuous measurement assay results

**Supplementary Figure 9** | Deconvoluted MS-1 plot for BV loading determination

**Supplementary Table 1**| Single-codon mutations screening and re-phenotyping

**Supplementary Table 2**| Wildtype smURFP and synHO coding sequences

**Supplementary Figure 1 |** Schematics of genetic circuits

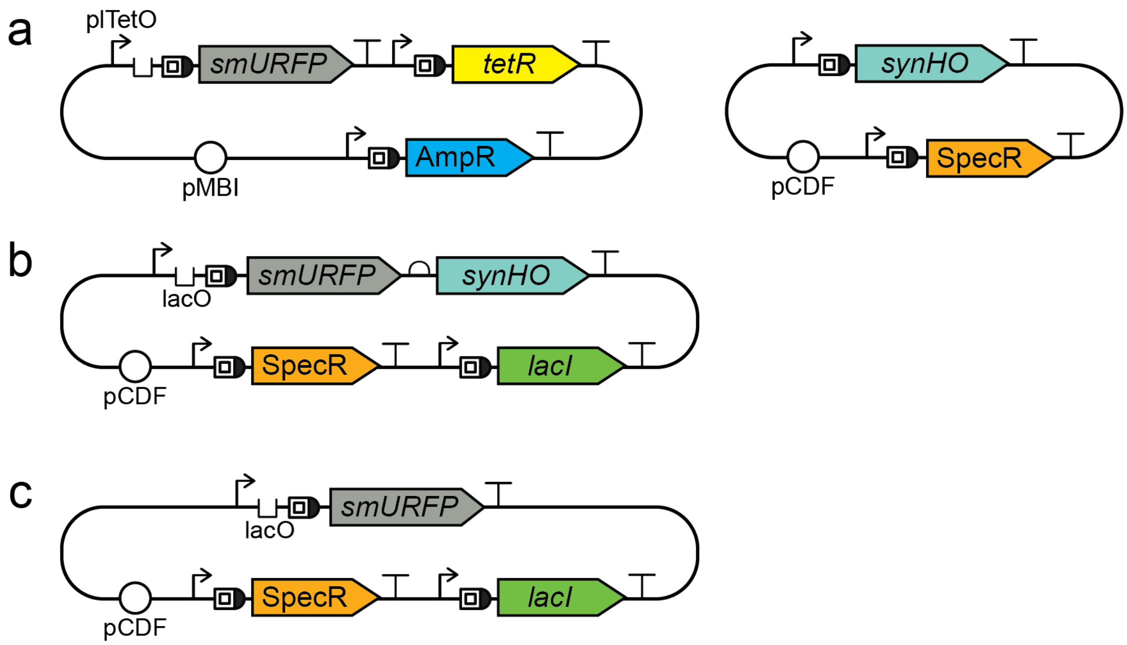


**Figure S1.** Cartoon schematics of the genetic circuits used to express smURFP. (a) The two-plasmid genetic circuit used for constructing and expressing the smURFP DMS library consists of a primary library plasmid and a secondary synHO plasmid. The primary plasmid encodes smURFP under the control of the plTetO promoter/operating and the cognate tetR repressor. The primary plasmid backbone features of an MBI origin of replication and the TEM-1 beta-lactamase resistance marker, labelled *AmpR*. The secondary plasmid encodes for constitutive expression of synHO and its backbone features a CDF original of replication. (b) pET plasmid with a CDF origin and spectinomycin resistance marker for expression of a bicistronic smURFP - synHO transcript under the control of LacO and a $P_{T7}$ promoter used for fermentation and purification of mature protein. (c) Alternative iteration of the pET plasmid for fermentation and purification of apo-protein.

**Supplementary Figure 2** | aTc induction gradients for the TetR-controlled circuit

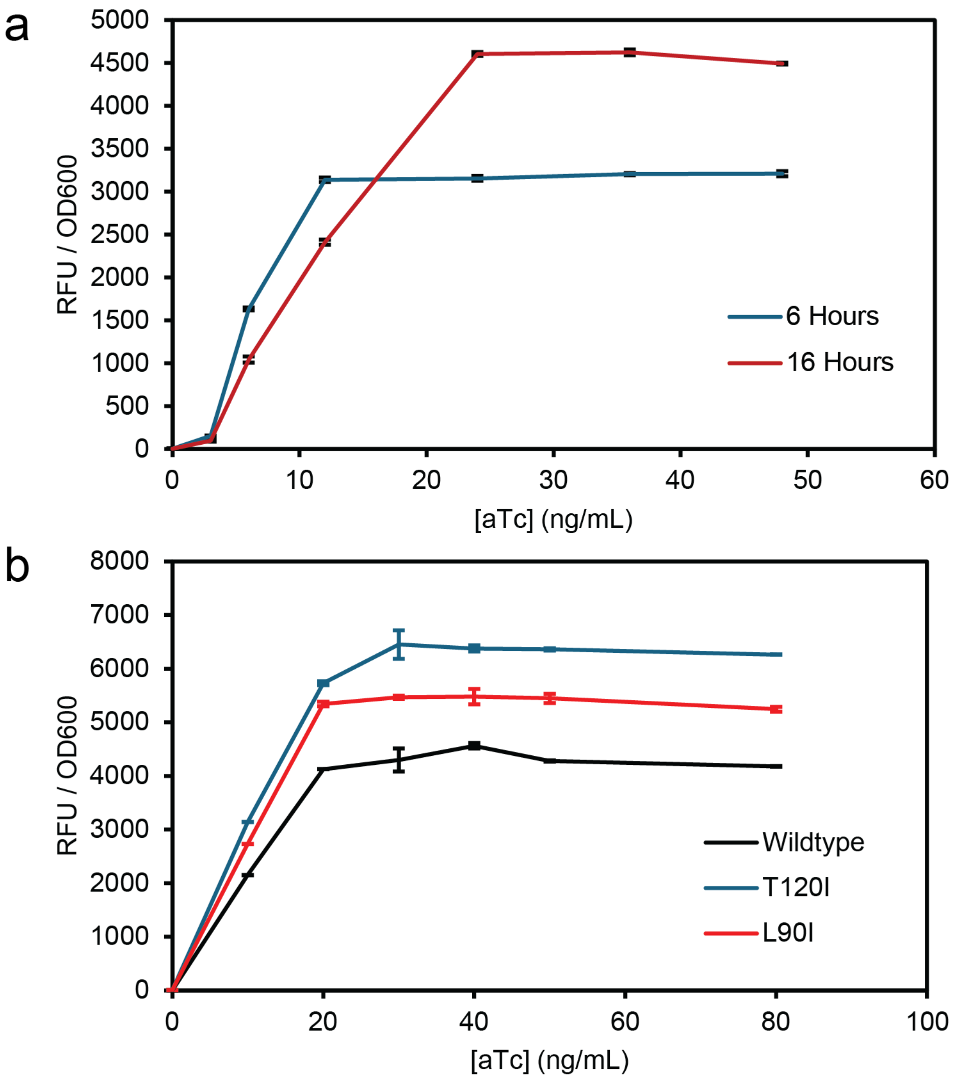


**Figure S2.** aTc induction gradient of TetR-controlled library circuit. (a) endpoint measurements of replicate samples at 6 hours and 16 hours. (b) induction curves for wildtype and two single mutants (T120I, L90I) measured at 16 hours. All measurements were performed in biological and technical triplicate and data represents mean fluorescence ± standard deviation, reported in relative fluorescence units (RFU) at an optical density of 600 nm.

**Supplementary Figure 3** | FACS-enriched library NGS coverage heat map

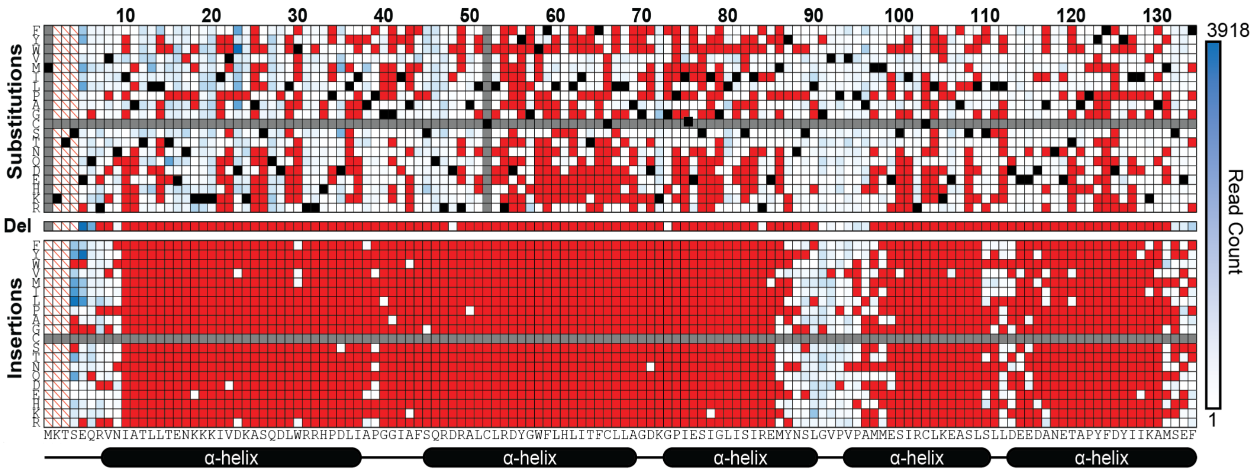


**Figure S3.** Heat map showing the direct read count coverage of all possible mutations in the FACS-enriched library with highlighted wildtype positions (black), unincluded mutations (gray), and unobserved expected sequences (red), and positions locked during amplification (white with red slash).

**Supplementary Figure 4** | Naïve versus enriched library singlicate screening example results

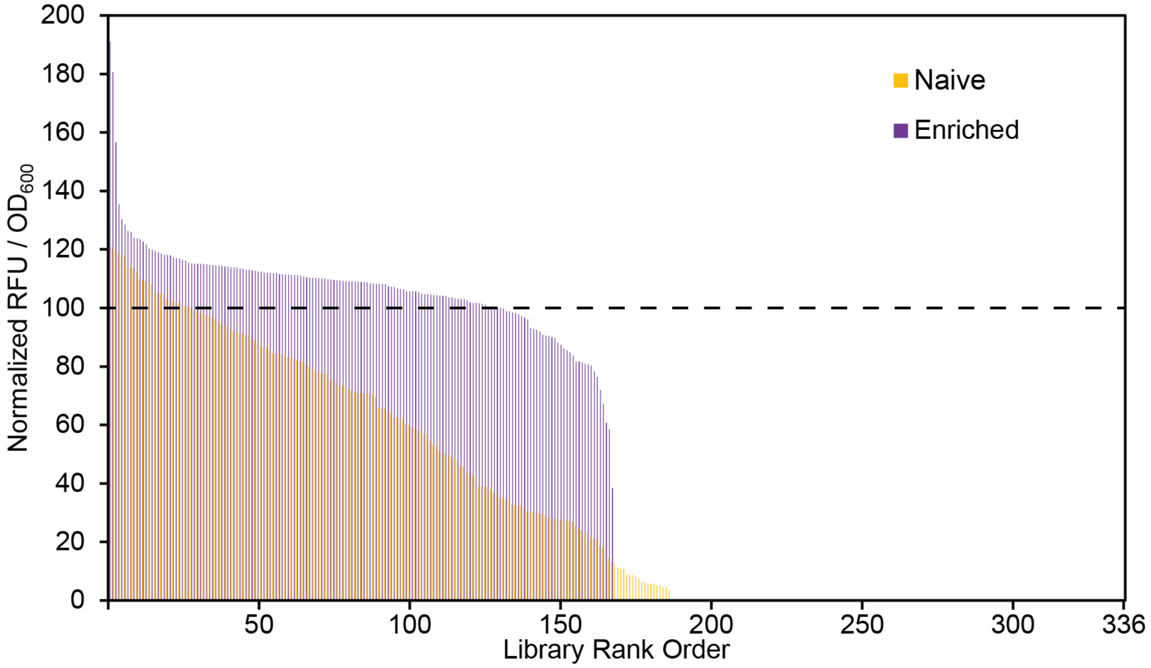


**Figure S4.** Singlicate fluorescence measurements of random clones from the naïve and enriched smURFP populations. Fluorescence is normalized to a wild-type control and represents 168 clonal isolates from the enriched library and 336 clonal isolates from the naïve library. Individual measurements from the two libraries are overlayed and the dashed black line indicated the wildtype signal.

**Supplementary Figure 5** | Absorbance and fluorescence spectra from *in vitro* measurement

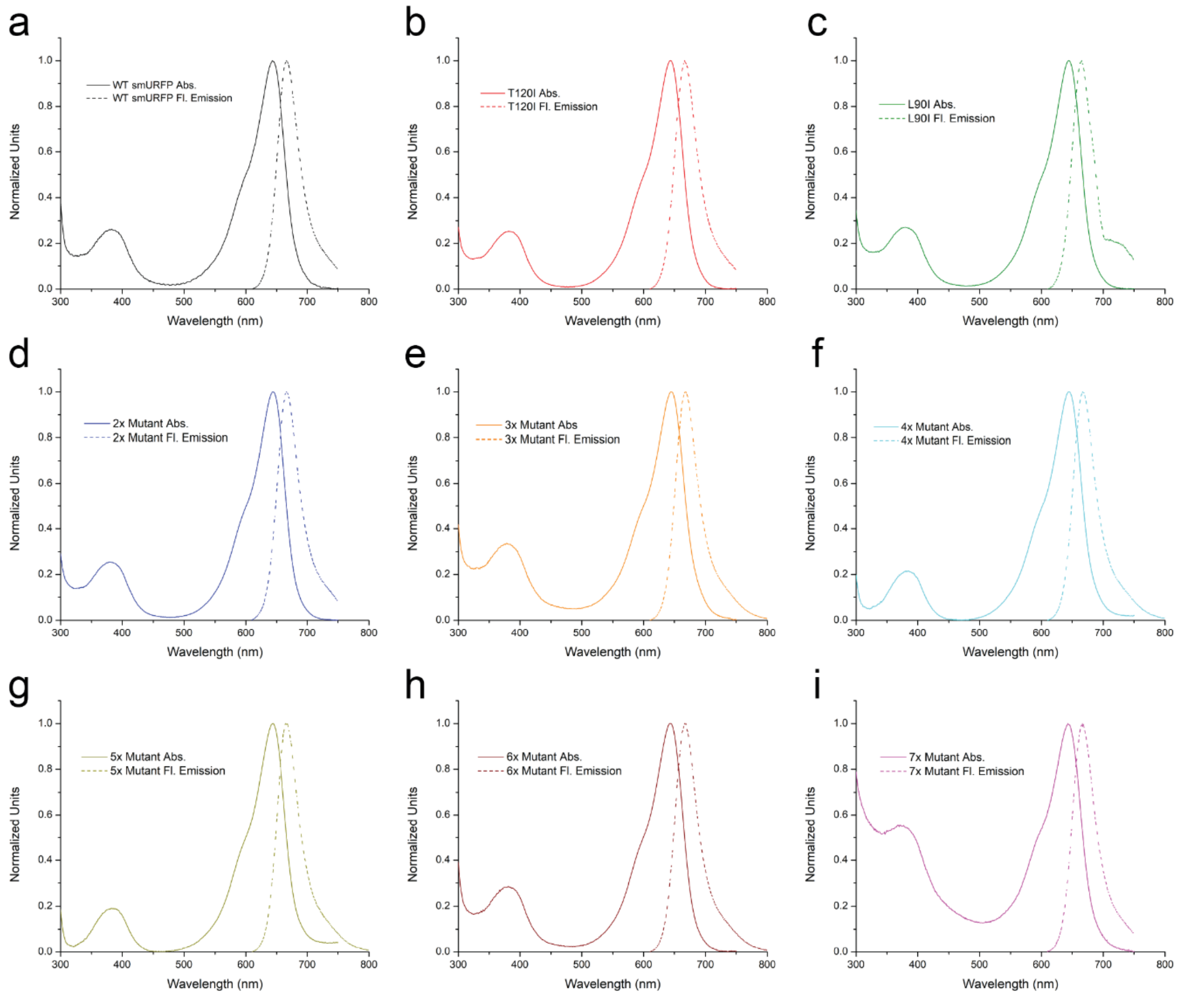


**Figure S5.** Normalized absorbance and fluorescence spectra from *in vitro* measurements for (a) smURFP wild type, (b) smURFP T120I (also denoted smURFP_2.1), (c) smURFP L90I, (d) smURFP_2.2, (e) smURFP_2.3, (f) smURFP_2.4, (g) smURFP_2.5, (h) smURFP_2.6, and (i) smURFP_2.7.

**Supplementary Figure 6** | Absorbance measurements for extinction coefficient calculation

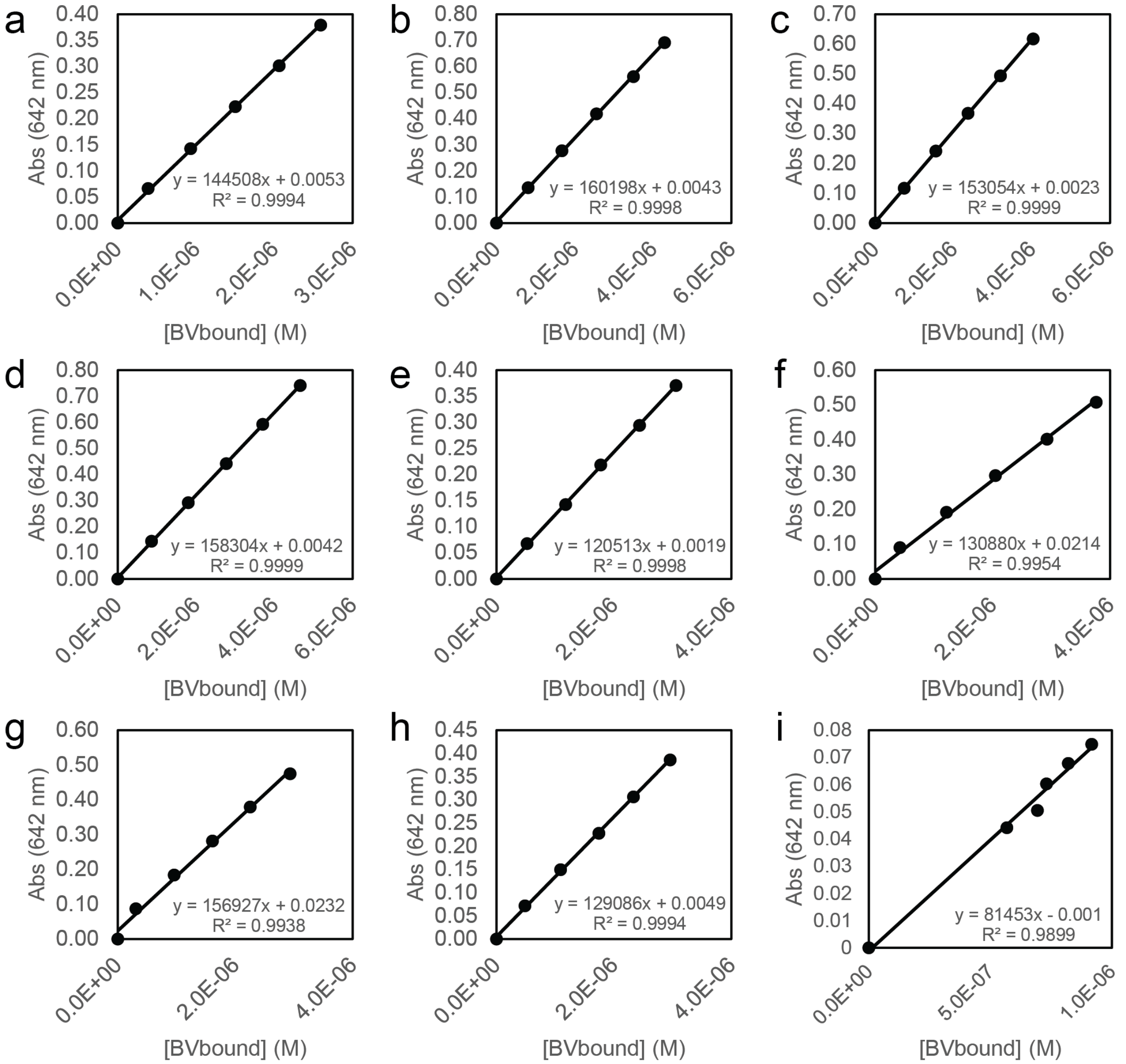


**Figure S6.** Calculation of molar absorptivity for FPs. Concentration of biliverdin bound was determined by Beer-Lambert-Bouguer's law at Soret band absorbance (391 nm; 39,900 $M^{-1}$ $cm^{-1}$) according to method described for (a) smURFP wild type, (b) smURFP T120I (also denoted smURFP_2.1), (c) smURFP L90I, (d) smURFP_2.2, (e) smURFP_2.3, (f) smURFP_2.4, (g) smURFP_2.5, (h) smURFP_2.6, and (i) smURFP_2.7.

**Supplementary Figure 7** | smURFP_2.1-2.7 phenotype under varied assay conditions

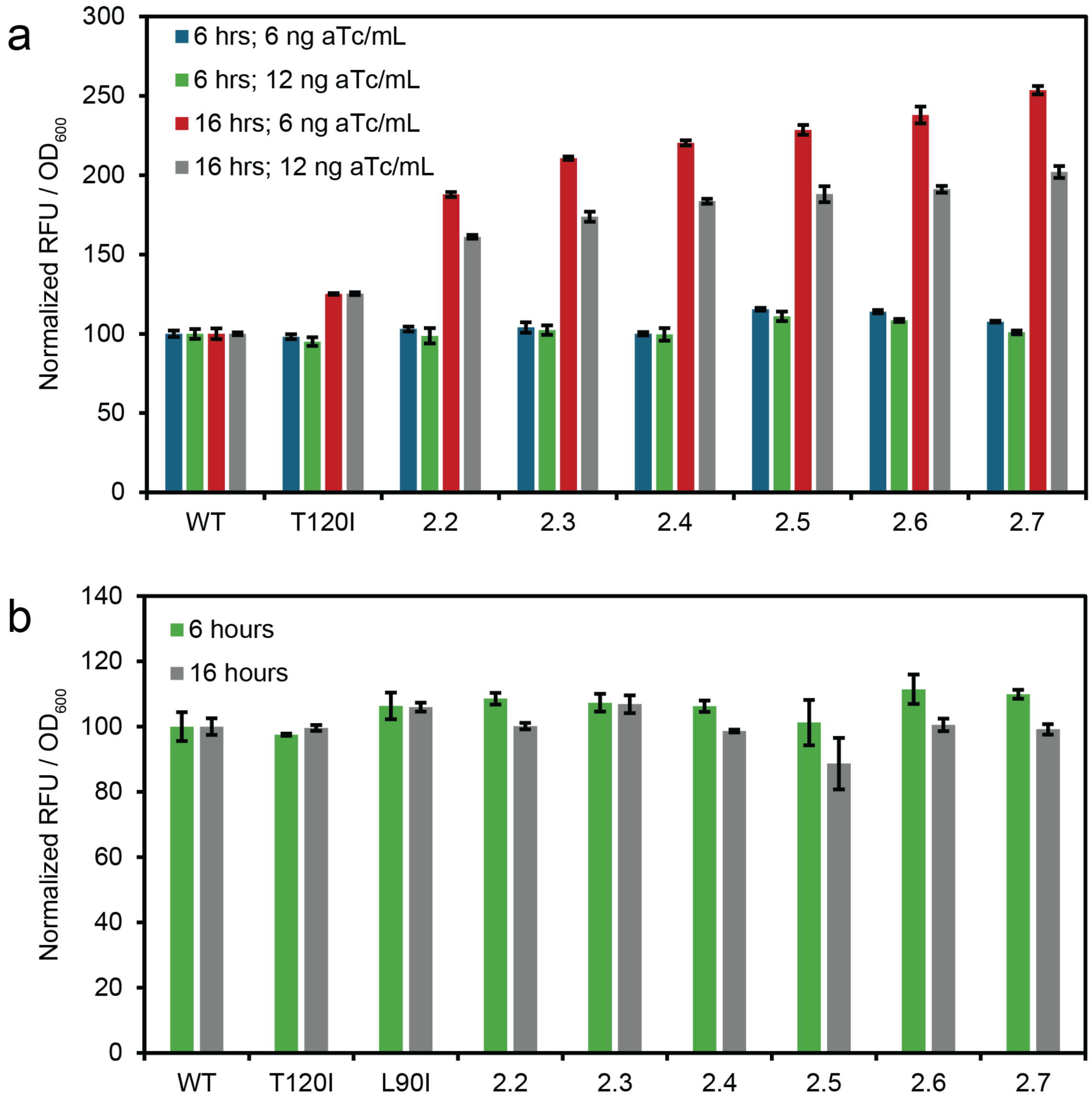


**Figure S7.** Influence of expression strength and timing on smURFP fluorescence during screening. End-point relative fluorescence of smURFP variants normalized to the wildtype for (a) the TetR-regulated plasmid system and (b) the bicistronic pET plasmid context. Data was collected in biological and technical triplicate and represents mean fluorescence ± standard deviation.

**Supplementary Figure 8** | Continuous measurement assay results

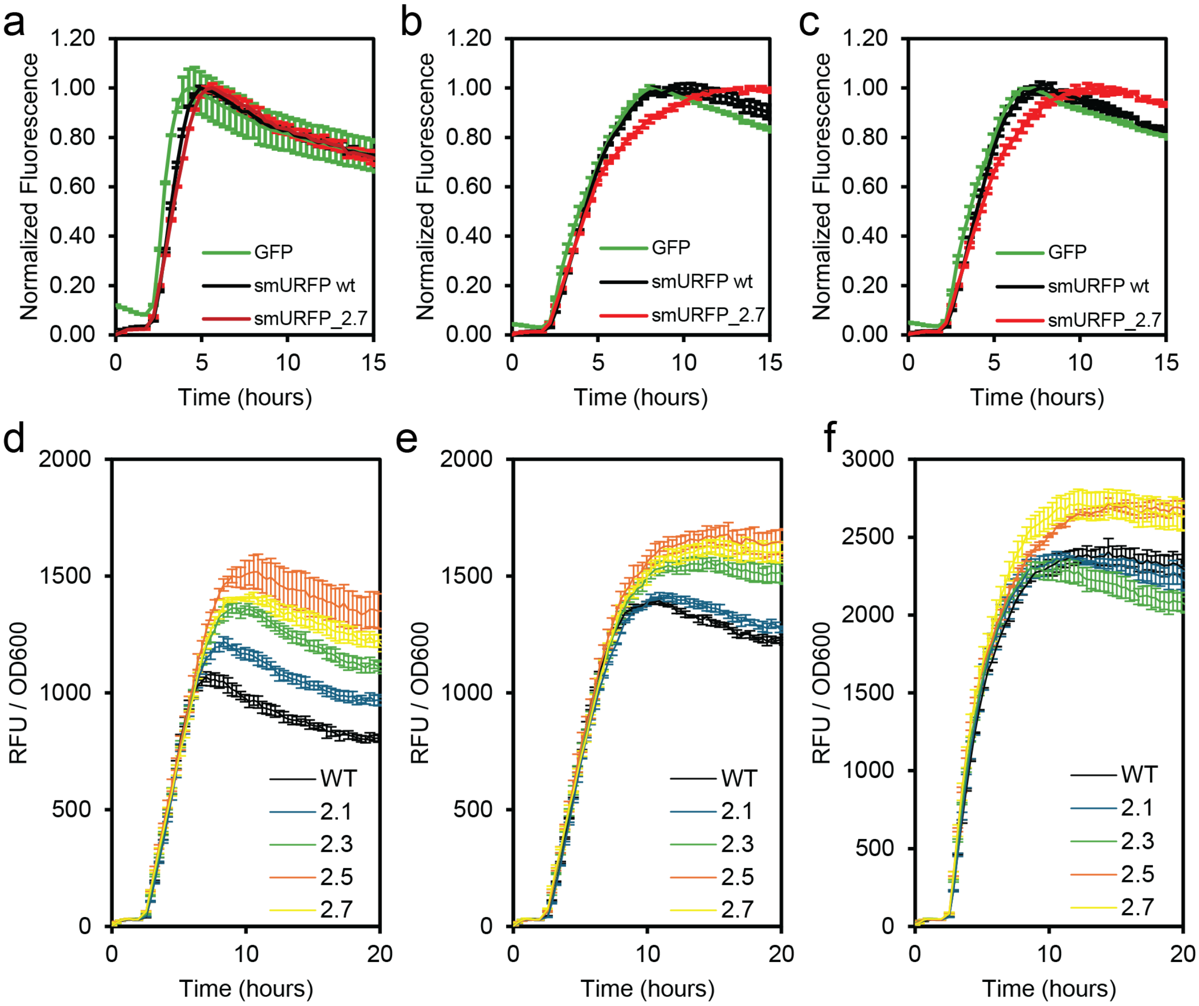


**Figure S8.** Time course measurement of RFU/$OD_{600}$ (normalized to the maximum average value per condition) for GFP, smURFP wild-type, and smURFP_2.7 in the TetR-controlled expression context induced with (a) 6 ng aTc/ mL, (b) 12 ng aTc/ mL, and (c) 24 ng aTc/ mL. Time course measurement of RFU/$OD_{600}$ for smURFP wild-type, 2.1, 2.3, 2.5, and 2.7 in the TetR-controlled expression context induced with (d) 6 ng aTc/ mL or (e) 12 ng aTc/ mL and (f) in the bicistronic pET expression context induced with 25 μM IPTG. All measurements were taken in biological triplicate and technical duplicate.

**Supplementary Figure 9** | Deconvoluted MS1 spectra for BV loading determination

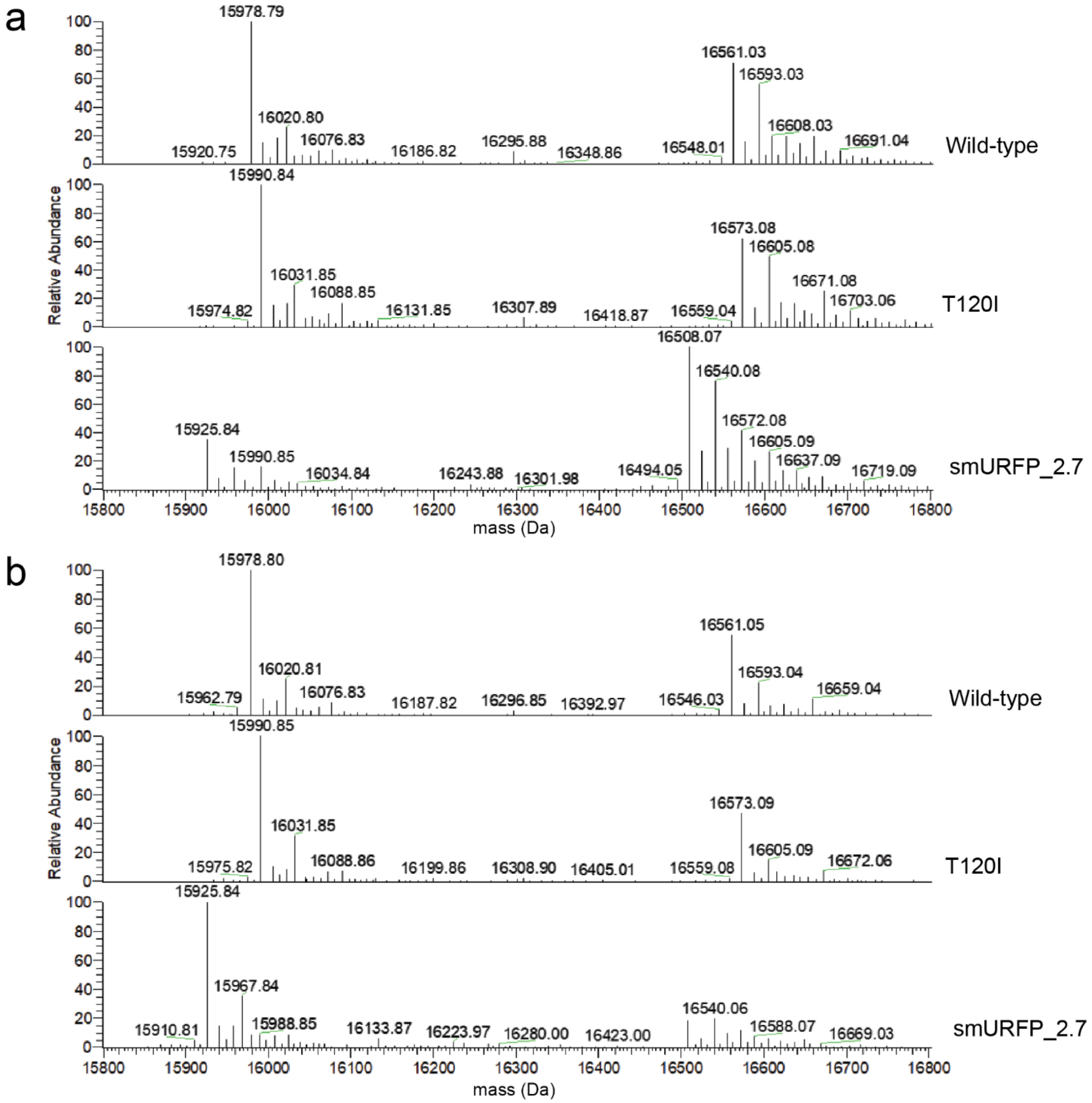


**Figure S9.** Expanded sections of deconvoluted mass spectra for smURFP wild-type, T120I, and 2.7 samples purified following aTc induced expression with (a) 6 ng aTc/ mL or (b) 12 ng aTc/ mL. The apo-protein and BV ligated masses are 15,978.79 Da and 16,561.04 Da, respectively, for wild-type, 15,990.84 Da and 16,573.09 Da, respectively, for smURFP T120I, and 15,925.82 Da and 16,508.06 Da, respectively, for smURFP_2.7. The abundances of the ion peaks corresponding to the apo-protein and BV-ligated protein were used to determine loading amounts.

**Supplementary Table 1|** Single-codon mutation screening and re-phenotyping

| Position | Mutation | Screened Plasmids | | Subcloned Plasmids | |
|---|---|---|---|---|---|
| | | Normalized RFU/$OD_{600}$ | Normalized StDev | Normalized RFU/$OD_{600}$ | Normalized StDev |
| 3 | K2 T3 ins M | 113.1 | 0.3 | - | - |
| 3 | T3M | 109.0 | 1.1 | - | - |
| 7 | Q6 R7 ins K | 115.6 | 1.9 | 110.3 | 2.3 |
| 10 | I10V | 106.4 | 0.2 | - | - |
| 12 | T12V | 108.3 | 1.6 | 112.2 | 1.7 |
| 12 | T12A | 115.8 | 0.4 | 105.8 | 2.8 |
| 13 | L13H | 105.1 | 0.3 | - | - |
| 20 | K20S | 116.0 | 0.8 | 103.7 | 4.0 |
| 27 | Q27H | 114.7 | 1.3 | 107.2 | 3.8 |
| 47 | R47A | 107.5 | 0.7 | 108.7 | 4.6 |
| 85 | E85Q | 122.3 | 1.0 | 113.4 | 2.7 |
| 90 | Del L90 | 127.2 | 0.2 | 124.0 | 2.3 |
| 90 | L90 G91 ins S | 136.8 | 2.0 | 130.2 | 2.1 |
| 90 | L90I | 108.1 | 0.7 | 154.3 | 4.1 |
| 90 | L90V | 141.6 | 1.5 | 102.8 | 1.1 |
| 91 | G91D | 112.8 | 1.0 | 114.0 | 4.1 |
| 91 | G91 V92 ins P | 117.3 | 0.9 | 126.4 | 4.0 |
| 91 | G91P | 120.0 | 1.1 | 126.2 | 1.6 |
| 92 | V92T | 106.4 | 0.4 | - | - |
| 93 | P93Q | 110.7 | 0.9 | 111.2 | 1.5 |
| 95 | P95G | 114.6 | 1.3 | 109.9 | 4.8 |
| 98 | M98Q | 108.6 | 2.2 | 103.9 | 1.4 |
| 118 | N118Q | 118.8 | 1.2 | 114.2 | 10.7 |
| 120 | T120I | 108.2 | 0.9 | 146.2 | 3.8 |
| 126 | Y126M | 115.8 | 1.3 | 103.2 | 5.5 |
| 132 | S132N | 115.6 | 1.7 | 113.7 | 1.6 |

**Supplementary Table 2**| Wildtype coding sequences for smURFP and synHO

| Protein | Coding Sequence |
|---|---|
| **smURFP (Wildtype)** | **ATG**aaaacttctgaacaacgtgtaaacatcgcaactctgctgactgagaacaaaaaga<br>aaatcgttgataaggcctctcaggatctgtggcgtcgtcatccagatctcattgcgcc<br>gggcggcatcgcttttagtcaacgcgatcgtgcactgtgcctgcgtgactacggctgg<br>tttctgcacctgatcacgttctgtctgctggctggtgataaaggtcctattgaatcca<br>tcggtctaattagcatacgcgaaatgtacaacagcctgggtgtgccggttccggctat<br>gatggaatctatccgttgtctgaaagaggcctccctgtcactgctggacgaggaagac<br>gccaatgagactgcaccgtactttgactatatcatcaaggctatgtccgaattccatc<br>atcaccatcaccat**TGA** |
| ***Synechocystis* Heme Oxygenase-1** | **ATG**agtgtcaacttagcttcccagttgcgggaagggacgaaaaaatcccactccatgg<br>cggagaacgtcggctttgtcaaatgcttcctcaagggcgttgtcgagaaaaattccta<br>ccgtaagctggttggcaatctctactttgtctacagtgccatggaagaggaaatggca<br>aaatttaaggaccatcccatcctcagccacatttacttccccgaactcaaccgcaaac<br>aaagcctagagcaagacctgcaattctattacggctccaactggcggcaagaagtgaa<br>aatttctgccgctggccaagcctatgtggaccgagtccggcaagtggccgctacggcc<br>cctgaattgttggtggcccattcctacacccgttacctggggggatctttccggcggtc<br>aaattctcaagaaaattgcccaaaatgccatgaatctccacgatggtggcacagcttt<br>ctatgaatttgccgacattgatgacgaaaaggcttttaaaaatacctaccgtcaagct<br>atgaatgatctgcccattgaccaagccaccgccgaacggattgtggatgaagccaatg<br>acgcctttgccatgaacatgaaaatgttcaacgaacttgaaggcaacctgatcaaggc<br>gatcggcattatggtgttcaacagcctcacccgtcgccgcagtcaaggcagcaccgaa<br>gttggcctcgccacctccgaaggc**TAA** |
